# A standardized, open-source, portable model for noninvasive joint injury in mice

**DOI:** 10.1101/2025.03.11.642661

**Authors:** Michael D. Newton, Lindsey Lammlin, Sofia Gonzalez-Nolde, Scarlet Howser, Isabelle Smith, Luke Stasikelis, Alexander J. Knights, RE-JOIN Consortium Investigators, Tristan Maerz

## Abstract

Preclinical models of osteoarthritis (OA) are crucial for the study of disease mechanisms and for the development of critically-needed disease-modifying therapeutics. While surgical OA models, such as the destabilization of the medial meniscus (DMM), have been the gold standard in the field for decades, noninvasive joint loading-based models have increased in popularity and utility. To facilitate standardization of the noninvasive anterior cruciate ligament rupture (ACLR) model in mice, we present the **Mo**bile **J**oint-Injury **O**perator **(MoJO)** - an open-source protocol with accompanying fixtures and data, designed for a low-cost, commercially-available, portable uniaxial testing system with a small footprint. We provide 3d-printable fixture designs and a rapid, highly-repeatable ACLR-mediated joint injury protocol that results in the expected post-traumatic osteoarthritis phenotype in male and female C56Bl/6 mice. We then describe the expected mechanical data from the injury procedure and offer various troubleshooting strategies. Finally, we summarize the resultant PTOA phenotype by knee hyperalgesia testing, µCT imaging, flow cytometry, and histological assessment. Increased standardization of this model is a critical aspect of the overall refinement of animal models of OA.

## Introduction

Preclinical animal models of osteoarthritis (OA) are invaluable for the study of disease mechanisms and the early development of novel disease-modifying therapeutics. While rats, guinea pigs, dogs, sheep, pigs, goats, and horses are also employed in clinical translation pipelines, mouse models remain the most widely utilized and most economical tool for preclinical investigation.

Among the strongest risk factors for the development of OA is joint injury, such as anterior cruciate ligament rupture (ACLR). Following ACLR, long-term human clinical studies demonstrate a ∼40% risk of developing post-traumatic OA (PTOA)^1^, with some studies indicating as high as 80%^2^. Given that the majority of human OA data has been collected at timepoints of already established disease, the early injury-induced mechanisms that initiate disease progression represent a major knowledge gap. Surgical OA mouse models are limited in their ability to assess early post-injury mechanisms given the need for surgical arthrotomy in the sham control group, which exhibits synovitis and hyperalgesia for up to 2-4 weeks post-injury^3, 4^.

Noninvasive joint injury models have become increasingly popular, and they complement surgical models in their utility to assess early post-injury processes. The tibial compression overload ACLR model (**Fig 1A**) is among the most widely utilized noninvasive model of PTOA in mice^5-8^ and rats^9-11^, and multiple variations have been developed, each with differences in equipment, fixtures, animal positioning, and loading protocols, resulting in a range of phenotypes. Subcritical cyclic loading has also been employed as a PTOA model with similar experimental setups^12, 13^. A relatively large barrier to entry associated with noninvasive models is the need for a mechanical testing system with high displacement and load resolution. These systems are generally costly ($100,000+) and require expertise to calibrate, tune, and operate. Due to their size and/or sensitive components, they are generally immovable, greatly restricting experimental flexibility (e.g. movement in and out of vivaria), especially in multi-institutional collaborations.

**Figure 1.**
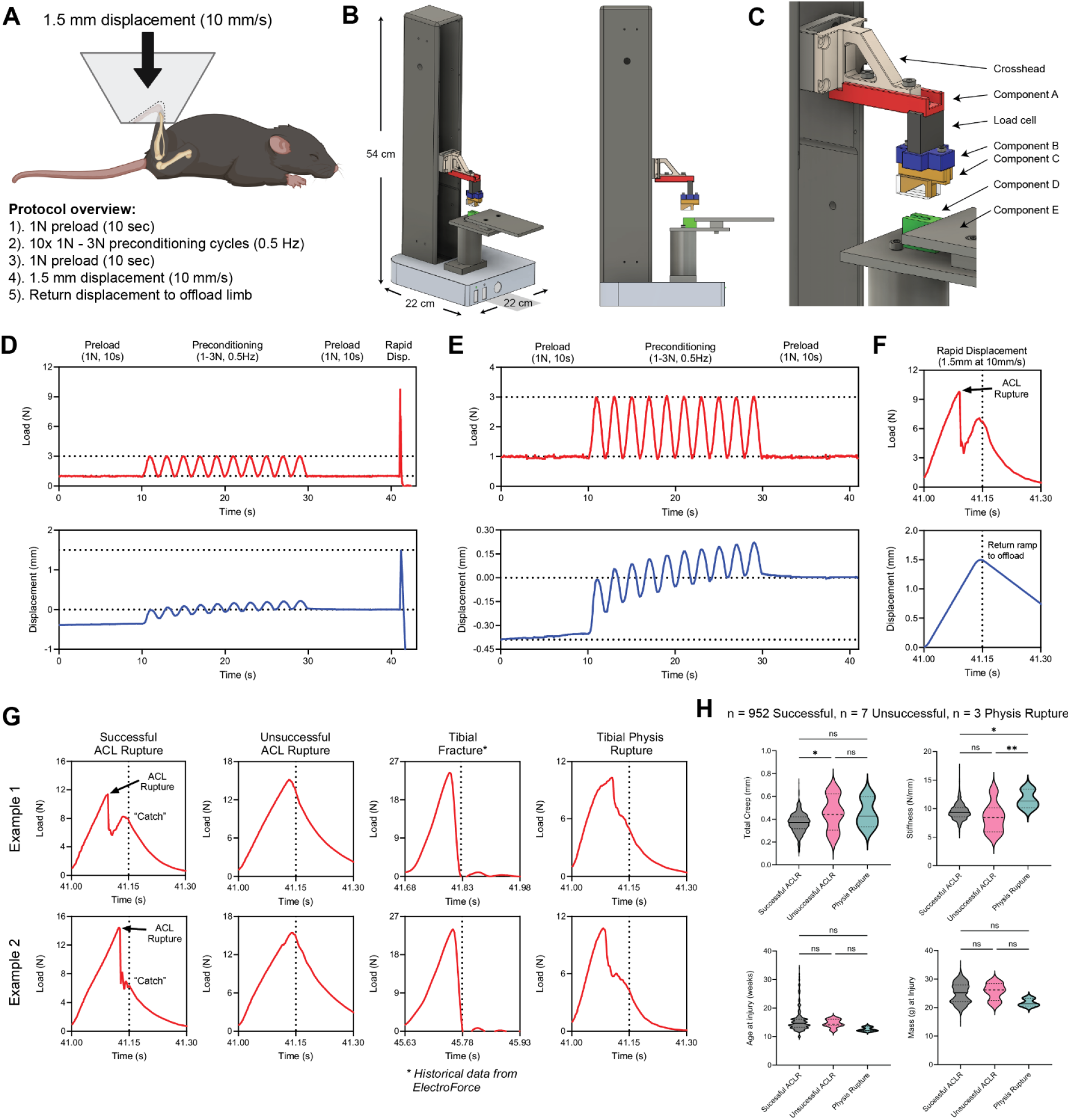
A). Schematic of mouse positioning and protocol overview. B). Isometric and side view of the Univert S2 and assembled ACLR fixtures. C). Isometric view of ACLR fixture components. D). Load vs time (top) and displacement vs time (bottom) plots of full loading protocol. E). Isolated view of load vs time (top) and displacement vs time (bottom) plots of preload and preconditioning segments to demonstrate expected creep. F). Isolated view of load vs time (top) and displacement vs time (bottom) plots of the final compressive loading segment to induce ACLR. The point of ACL failure is noted. G). Two representative examples of load vs time plots of successful and unsuccessful ACL ruptures, tibial fractures, and tibial physis ruptures. Note the characteristic secondary “catch” that should be observed following ACL rupture. H). Quantitative comparison of mechanical data, age at injury, and mass at injury between mice in successful ACLR, unsuccessful ACLR, and physis ruptures.

To facilitate standardization and ease of access to the ACLR model in the preclinical OA community, we present an open-source protocol with accompanying fixtures and data, designed for a low-cost, commercially-available, portable uniaxial testing system. The **Mo**bile **J**oint-Injury **O**perator **(MoJO)** demonstrates a high degree of repeatability, and hyperalgesia phenotyping, cellular analyses, and histological assessments confirm the expected PTOA phenotype consistent with our prior studies employing a conventional, immobile mechanical testing system.

## Methods

### Mechanical Testing System and Load Cell

Our protocol utilizes the Univert S2 (CellScale Inc., Waterloo, ON, Canada), a lightweight (8 kg), low footprint (22×22×54 cm), portable electromechanical testing system. However, the injury protocol and associated fixtures may be adapted to any given mechanical testing system with performance specifications similar to those of the CellScale Univert S2 (see manufacturer specifications). Given the need for high acceleration and displacement rate over a short displacement, electromechanical systems are recommended, as servohydraulic systems are generally less capable of achieving target velocities over the short displacements used for mice. During the rapid compressive ramp in our protocol, the Univert S2 crosshead accelerates at a consistent 650 mm/s^2^, reaching the target 10 mm/s velocity in 14.3 ms (∼9.5% of the total 150 ms displacement event) and over a 0.1 mm distance (∼6.7% of the total distance). After reaching the target displacement, the crosshead decelerates at 604.5 mm/s^2^, reaching 0 mm/s in 18.7 ms (12.5% of the total 150 ms displacement event) and over a 0.1 mm distance (∼6.7% of the total distance).

For the utilization of this protocol in mice, a 20 N or 25 N load cell is recommended. This ensures accuracy in the preload and preconditioning segments while ensuring that the ultimate injury load, generally 9-15N in C57BL/6 mice, does not exceed the maximum load specification. The load cell utilized for our design is the Zhimin ZMSA 2 kg S beam load cell (Anhui Zhimin Electrical Technology Co., Ltd., Bengbu, China), which has a single M3 mounting screw at each end for assembly.

### Fixtures

Our fixture design (**Fig. 1B-C, Suppl. Fig 1A-C**) is comprised of a minimal list of components, all amenable to 3d printing, with standard M3 and M5 metric fasteners used for assembly. To minimize the chance for fixture deflection/deformation and to maximize fixture lifespan, we recommend a high-stiffness, high-toughness plastic for 3d printing. We employ Tough 2000 resin, printed on a Form 3B printer (Formlabs Inc., Somerville, MA, USA). The 3D designs for all components are available on the associated Github repository (https://github.com/um-maerz-lab/MOJO/tree/main/), including both .STL files as well as component and assembly files for Autodesk Fusion360 (available at no cost to all educational institutions via autodesk.com/products/fusion-360/, to date). **Suppl. Fig. 1** is a detailed component and assembly plan. In our implementation, the animal bed assembly (components E1-4) was machined out of aluminum stock, but our design files include models for 3D printing these parts. Additional details on fixture design and adaptation to different setups are provided in **Suppl. Methods 1**.

### Mouse Positioning and Procedure

Following anesthetic induction, mice are placed prone on the animal bed (E1) and the knee of either hindlimb is placed over the cranial edge of the knee fixture (D) (**Suppl. Fig 1D**). The hindpaw is then gently manipulated into Component C, visualizing complete seating of the hindpaw into the fixture. This is most easily accomplished with two operators, one mouse handler to simultaneously guide the hindpaw and knee into their respective troughs, and a second computer operator who slowly lowers the crosshead until a low load (<0.2-0.5 N) is registered. Limb manipulation is easily performed with small, blunt manipulators (e.g. forceps), with one holding the knee in the trough while the other guides the hindpaw into the hindpaw fixture (C). By gently applying traction to the mouse in the cranial direction, secure positioning of the knee in the knee trough (D) is ensured, which is confirmed via gentle medial-lateral manipulation. Appropriate mouse positioning is illustrated in **Suppl. Fig. 1D**.

Mice tolerate this procedure well. Healthy mice do not exhibit any post-injury weight loss or alterations to general-health indicators. Inhaled isoflurane is sufficient for anesthesia (4-5% for induction, 1-2% for maintenance, flow rate 0.4-0.8 liter/min). Post-procedural analgesia may vary study-by-study, and we employ both carprofen (a single 5 mg/kg subcutaneous injection) or buprenorphine XR (a single 3.25 mg/kg subcutaneous injection).

Following positioning, a downward, compressive load is manually applied to the joint until ∼1 N is reached, and the automated protocol is initiated. The injury loading protocol involves five steps (**Fig. 1D**):

1. A 1N preload, held for 10 seconds (load controlled)
2. Ten sinusoidal preconditioning cycles between 1-3N, 0.5 Hz (load controlled)
3. A second 1N preload for 10 seconds (load controlled)
4. The injury-inducing 1.5 mm downward displacement, 10 mm/s (displacement controlled)
5. A 5 mm upward return displacement to offload the hindlimb, 5 mm/s (displacement controlled)

### Determination of Successful ACL Rupture

Examples of mechanical outputs from successful and unsuccessful ACL rupture procedures are shown in **Fig. 1G**. Success of the ACL rupture procedure can be reliably assessed via a combination of mechanical data assessment and visual/auditory cues. Successful ACL rupture will result in an abrupt drop in load as the ACL fails and the tibia subluxes anteriorly, but *this should not result in full unloading of the joint* – a “catch” and/or secondary re-loading will generally be observed as downward displacement continues and other joint tissues (namely meniscus) provide resistance. For larger mice, which undergo ACL rupture later in the 1.5-mm displacement stage, re-loading may not occur, but the “catch” should still be observable (**Fig. 1G**, Example 2). A successful ACL rupture should produce a characteristic audible “pop” that can be recognized well with repetition. Anterior subluxation of the tibia can also be visually appreciated, though the event is very rapid. Successful ACL rupture can be physically assessed via the anterior drawer test^9^, though in mice this procedure is challenging to interpret. When first adopting the model, successful isolated ACL rupture should be confirmed via *postmortem* dissection in trial cohorts. In our hands, the success rate of this procedure is incredibly high (99.0%), but unsuccessful procedures do occur. In our experience, unsuccessful ruptures produced characteristically different loading profiles and audiovisual feedback (**Fig. 1G**). Additional details for recognizing and diagnosing both successful and unsuccessful ruptures are provided in **Suppl. Methods 2**.

### Mechanical Tester Tuning

Correct tuning is essential for load-controlled movements to execute accurately. The Univert S2 is equipped with simplified, pre-programmed velocity and acceleration tuning (scale 0-10), and we have found that values of 9 for both parameters produce excellent results. These parameters serve as simplified alternatives to full proportional– integral–derivative (PID) tuning, though advanced options do allow for further fine-tuning of PID values if desired. A protocol file for the Univert S2 control software is available on the Github repository. The tuning parameters we employ are embedded with this file. If adjustments to tuning are desired, we recommend using a steel compression spring – this can also be used to confirm correct performance at the start of each testing day, prior to live animal experimentation (see **Suppl. Methods 3** for more information).

***Caution:*** *Changing PID parameters can cause any mechanical testing system to become unstable and oscillate at high frequency. To avoid injury, only experienced operators should perform PID tuning, and a safety shield and safety glasses should always be used during tuning*.

### Mechanical Data Analysis

We developed a MATLAB code to automatically and unbiasedly analyze mechanical data outputs from this protocol. This script is available on the Github repository and has been annotated for clarity. Data outputs include rupture displacement, ultimate displacement, rupture load, ultimate load, linear elongation stiffness, percent error in preconditioning load amplitudes, and percent error in ultimate displacement. These data, in addition to a graphical output of displacement vs time and load vs time plots, are important quality control tools and should be reviewed following each procedure to confirm successful and repeatable ACLR. To permit prospective evaluation of protocol repeatability, this study’s Github repository contains a sample of mechanical data outputs from our MoJO procedures. An analogous code has also been provided to analyze the results of a test protocol run on the calibration spring, which we recommend using to monitor the performance of the testing system and fixtures over time.

To mechanically validate the MoJO system, we performed bilateral ACLRs in a cohort of mice (*n*=6) whereby one limb underwent ACLR using the MoJO system and the other limb underwent ACLR using our established ACLR platform on our conventional, immobile mechanical testing system and associated fixtures (ElectroForce 3300AT, TA Instruments, New Castle DE, USA)^8, 14, 15^. Analysis of mechanical outputs were performed in paired fashion between the two limbs.

### Flow Cytometry of ACLR-induced synovial inflammation

To confirm that the MoJO protocol recapitulates the injury-induced synovitis we have characterized in the ACLR model using the conventional ElectroForce system, male C56Bl/6 mice were subjected to bilateral ACLRs (i.e. one limb via MoJO, the other via ElectroForce)^8, 14, 15^. At 7 days post-injury, whole knee synovium (inclusive of Hoffa’s fat pad) was microdissected and digested enzymatically, as previously described^14, 15^, and flow cytometry was performed to assess major synovial cell populations (**Suppl. Methods 4)**.

### Confirmation of PTOA Phenotype

Male and female C57BL/6 mice aged 12-14 wks were used to demonstrate the behavioral, histological, and structural phenotype of the MoJO protocol. All mice underwent unilateral ACLR or an anesthesia/analgesia-only sham procedure, and were euthanized at 28 days post-injury, representing a timepoint of moderate to established disease severity. Inhaled isoflurane was used for procedural anesthesia, a single carprofen dose (5 mg/kg subcutaneous) was administered immediately after ACLR, euthanasia was performed via CO2 asphyxia, and hindlimbs were immediately dissected and fixed in 10% neutral-buffered formalin for 48 hrs and transferred to 70% ethanol for storage.

Blinded pressure application-based knee hyperalgesia testing was performed at baseline and 7, 14, and 28 days post-ACLR, as previously described^1^, using contralateral joints as controls. The average of triplicate measurements was used.

Hindlimbs were rehydrated in PBS overnight and imaged by µCT (8.9 µm voxel, 55 kVp, Bruker Skyscan). Limbs were paraffin processed, embedded, and sectioned in the sagittal plane (5 µm sections). Safranin-O/Fast-Green staining was used to demonstrate the expected phenotype in the medial and lateral compartments^14^.

### Statistical Analyses

Statistical comparisons were performed using Prism (v10.3.0, GraphPad Software LLC, San Diego, CA, USA) and SPSS (v29.0, IBM Corp., Armonk, NY, USA). Comparisons between MoJO and ElectroForce systems in matched mechanical, flow cytometry, and hyperalgesia data were performed via one-way repeated-measures ANOVA with system as the between-subject effect. Correlations of sex, age at injury, and mass at injury to mechanical parameters were performed via Pearson correlation analysis. Comparisons in mechanical data between operators and laboratories were assessed via ANCOVA, with operator or institution as the between-subject effect and age and sex as cofactors.

## Results

### Mechanical Data

Analysis of mechanical data from *n*=952 MoJO ACLRs demonstrates a high degree of success and repeatability. The overall rate of successful ACL ruptures was 99.0% (952 / 962). The ten unsuccessful procedures were comprised of 7 unsuccessful ACL ruptures and 3 tibial physeal displacements. Analysis of these cases shows no significant difference in age or body mass of the mice (**Fig. 1H**), but mechanical analysis shows that unsuccessful ACLRs had greater total creep during preloading and preconditioning, consistent with improper positioning (**Fig. 1H**). Physis ruptures had higher stiffness during linear displacement, also likely an indicator of poor positioning and engagement of the long bones rather than the ACL.

We observed very low error in reaching target loads during preconditioning and in reaching the target displacement of the compressive rupture ramp (**Fig. 2A**). The total displacement during the rupture event ranged 1.498-1.500 mm, never exceeding the target displacement. Failure displacement, failure load, stiffness during linear elongation, and total creep also exhibit high consistency, with expected differences between male and female mice (**Fig. 2B**). While the majority of our mice are C57BL/6 mice aged 12-14 weeks, our group also employs multiple transgenic models (most with a C57BL/6 background) that exhibit a range of skeletal sizes and phenotypes, but this analysis does not differentiate these. We observed some significant but very weak linear correlations between failure load/displacement and the body mass and age of mice (**Fig. 2C-D**).

**Figure 2.**
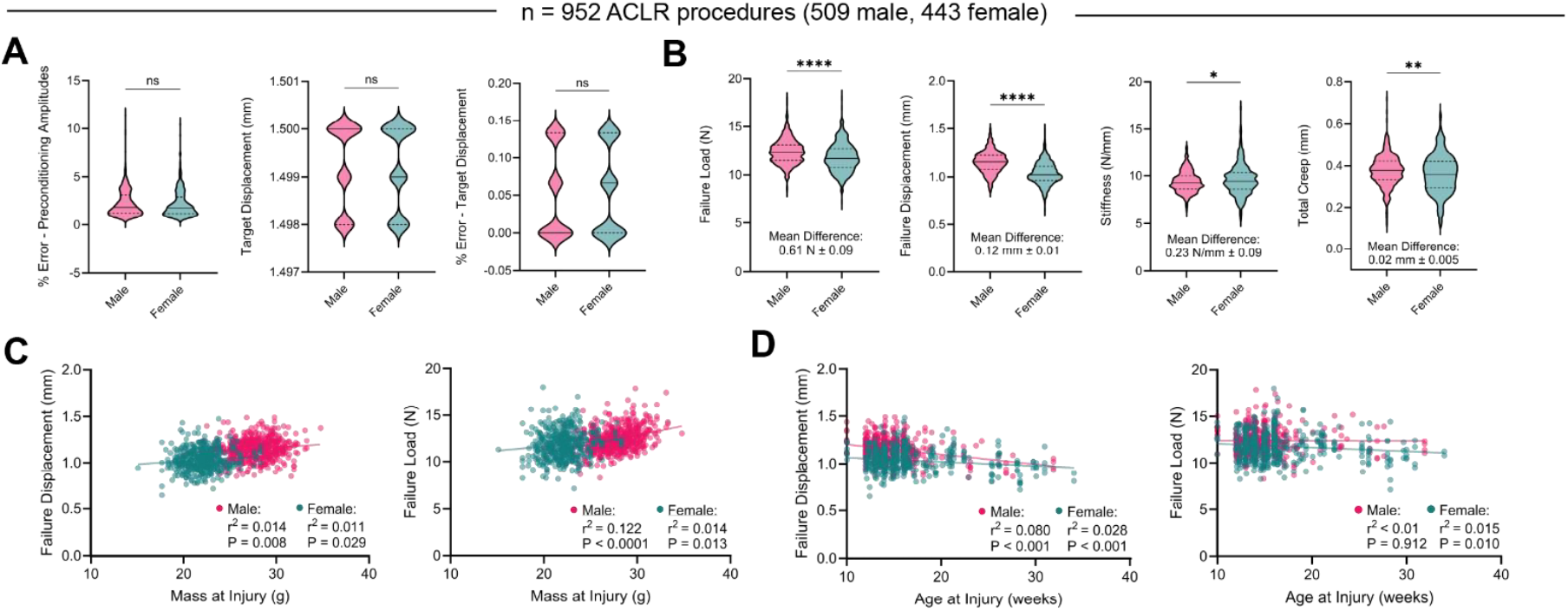
A). Accuracy and repeatability data of the ACLR protocol with the Univert S2. B). Descriptive ACLR outcomes derived from analysis of mechanical data, stratified by sex across n=509 male and n=443 female mice. C). Linear correlations between body mass at the time of injury and failure displacement (left) and failure load (right). D). Linear correlations between age at the time of injury and failure displacement (left) and failure load (right). P and r^2^ values were calculated via Pearson bivariate correlation analysis.

To interrogate repeatability between operators, we analyzed mechanical data between 4 experienced operators within our laboratory (*n*_*1*_=313, *n*_2_=289, *n*_3_=245, *n*_4_=108 ACLRs respectively). There were significant differences in mouse sex and age between operators, and these were statistically accounted for as cofactors (**Suppl. Fig. 3A**). Failure load, failure displacement, and stiffness were all highly similar between operators, with mean percent differences <3% for all parameters (**Suppl. Fig 3B**). Minor, though statistically significant differences were detected in failure load and stiffness between operators, likely due to small differences in limb positioning, coupled with high statistical power. We further compared mechanical data from our laboratory (*n*=955) to that of a collaborating laboratory (*n*=43) utilizing a separate but identical implementation of the MoJO system (**Suppl. Fig. 4A**). Again, we found comparable mechanical outcomes between institutions (**Suppl. Fig. 4B**), with only minor (though significant) differences in failure displacement and stiffness (**Suppl. Fig. 4C**).

**Figure 3.**
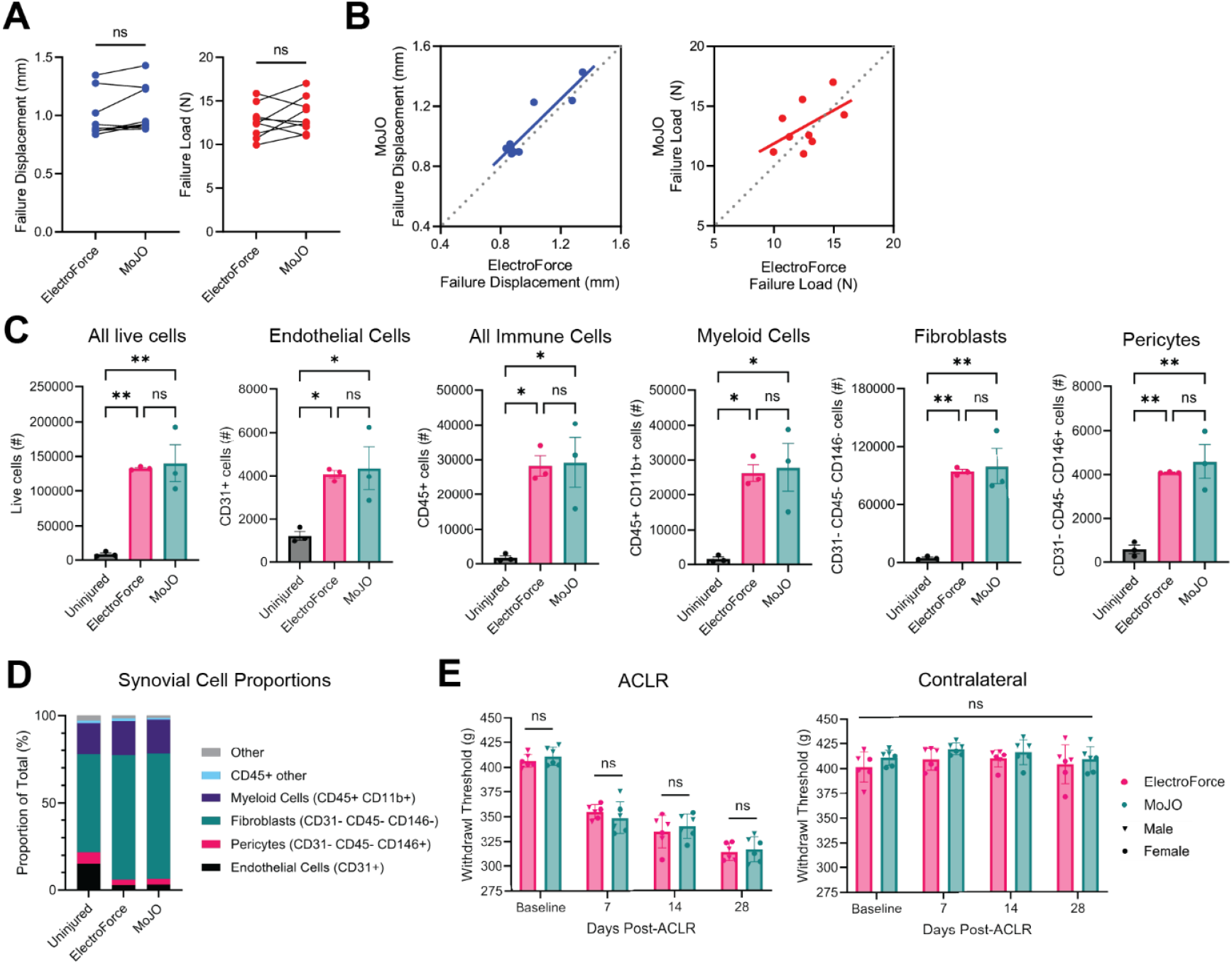
A). Failure displacement (left) and failure load (right) in a cohort of matched ACLR procedures in which the ElectroForce protocol was employed on one limb and the MoJO protocol on the other. B). Linear correlations between ElectroForce- and MoJO-derived failure displacement (left) and failure load (right). C). Flow cytometric assessment of synovial cells in naïve/uninjured mice, ElectroForce-injured mice, and MoJO-injured mice. n=3 samples per condition, each made up of two pooled synovia. D). Synovial cell proportions derived from flow cytometry between uninjured, ElectroForce-injured, and MoJO-injured mice. E). Knee withdrawal thresholds from ACLR joints (left) and contralateral joints (right), derived from blinded longitudinal knee hyperalgesia testing in ElectroForce-injured and MoJO-injured mice. n=6 mice per group assessed longitudinally.

**Figure 4.**
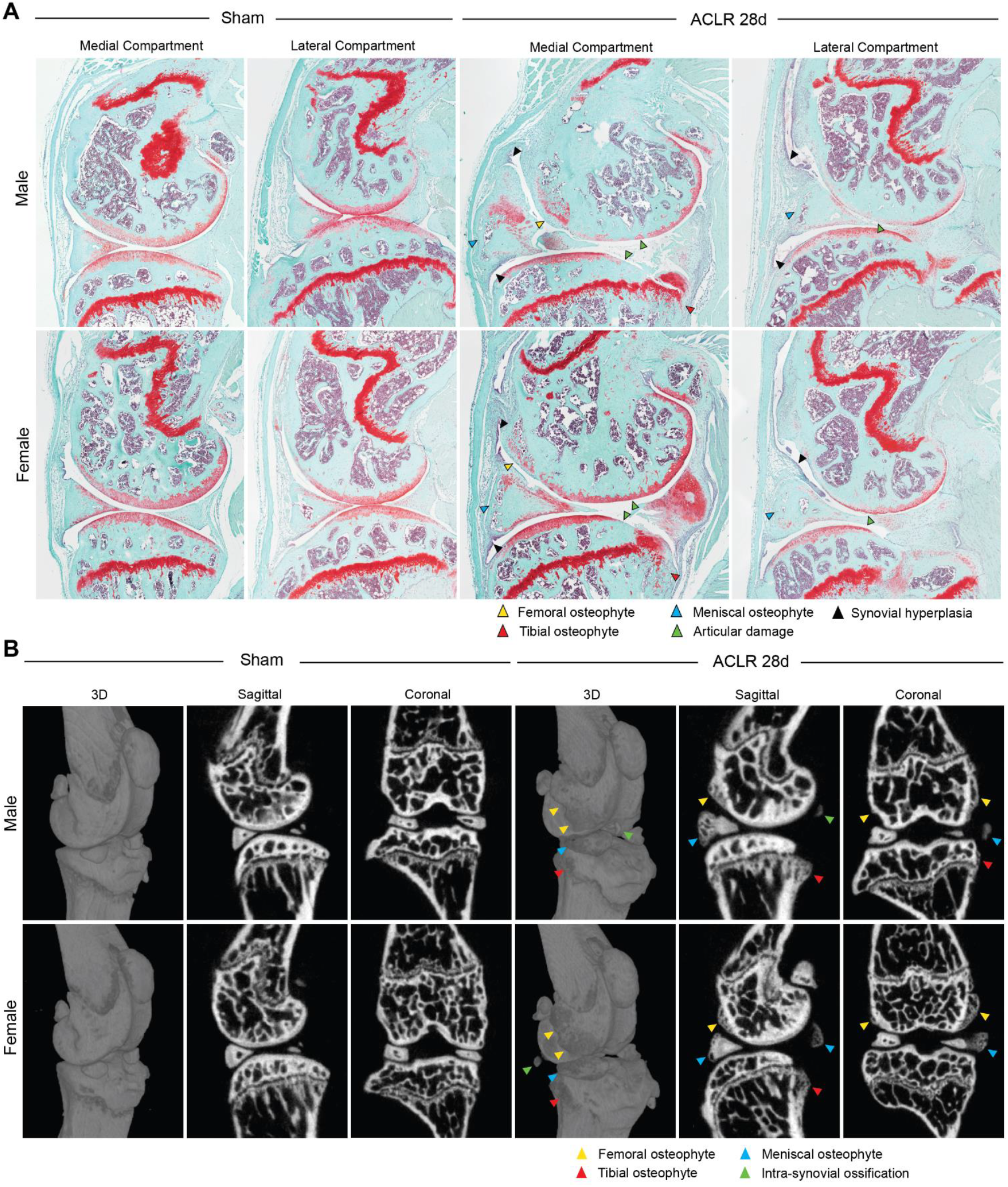
A). Safranin-O/Fast-green staining of sagittal sections from the medial and lateral compartments of Sham and ACLR 28d mice. B). 3D, sagittal, and coronal views of whole joints, derived from µCT imaging. Expected phenotypic characteristics are labeled with arrowheads.

These findings confirm the highly repeatable, reproducible, and accurate nature of this model. Nevertheless, each group should develop their own quality control processes with respect to acceptable mechanical outcomes, ideally corroborated by gross dissection to confirm ACL rupture and to rule out unexpected injuries.

### Comparison to ElectroForce system

In a comparative cohort of bilateral ACLRs, in which one limb underwent ACLR with the MoJO system and the other limb underwent ACLR with our conventional system using an ElectroForce mechanical tester, we found a high degree of repeatability between the two systems and protocols (**Fig. 3**), confirming that the MoJO protocol recapitulates our conventional protocol. There were no differences in failure displacement or failure load at the point of ACLR between matched MoJO-injured and ElectroForce-injured joints (**Fig. 3A**), and these outcomes were linearly correlated between the two limbs (**Fig. 3B**).

As a readout of ACLR-induced synovitis, the primary focus of our laboratory, we performed flow cytometric analysis of bilateral (i.e. MoJO on one limb vs ElectroForce on the other limb) synovial tissue 7 days following ACLR, employing sham mice as uninjured controls. We observed the expected increases in synovial endothelial, immune, and stromal cells in both ACLR groups, with no significant differences in absolute cell number (per single synovium) of live cells or any subset between the two models (**Fig. 3C**). The injury-induced shift in cell proportions between uninjured and ACLR synovia was highly similar between MoJO and ElectroForce (**Fig. 3D**). We have extensively characterized the phenotypic diversity of the immune^16^ and stromal cells^15^ that underpin this ACLR-induced synovitis.

To confirm that the two models induce similar alterations in pain behavior, we performed blinded longitudinal knee hyperalgesia testing in separate cohorts of age- and sex-matched mice injured unilaterally with the two systems. Following injury, we observed the expected reductions in knee withdrawal thresholds in ACLR joints, and there were no differences in withdrawal threshold at baseline or any post-ACLR timepoint between MoJO and ElectroForce (**Fig. 3E**).

Taken together, these data demonstrate that the MoJO system accurately recapitulates our conventional protocol using the ElectroForce system, with equivalent mechanical induction of the injury, similar alterations in synovial cell populations, and highly comparable onset of knee hyperalgesia.

### Structural PTOA Phenotype

To illustrate the expected PTOA phenotype using the MoJO system, we performed histological assessment and µCT imaging in Sham or ACLR 28d mice. Consistent with our prior sex-dependent characterization of this model using our conventional ElectroForce protocol^14^, we observed the expected onset of articular and meniscal damage, the formation of osteophytes and chondrophytes, most prominently at the anteromedial meniscus, and subchondral bone sclerosis (**Fig 4A**). Articular damage was most severe at the anterior aspect of the medial femoral condyle and the posterior aspect of the medial tibial plateau, where we observed complete erosion of noncalcified cartilage down to calcified cartilage, and in most severe examples, down to subchondral bone (**Fig 4A**). Articular damage is much milder in the lateral compartment, where we observed superficial articular damage but no massive lesions or complete erosion. Osteophyte formation is also milder in the lateral compartment.

The synovial phenotype was also consistent with our prior published work, characterized by synovial lining hyperplasia, subsynovial fibrosis and immune cell infiltrate, volumetric loss of Hoffa’s fat pad, and synovial exudate (**Fig 4A**). In the posterior synovium, focal cartilage formation was observed at varying degrees, and a synovial pannus was occasionally identified as invading and covering anterior tibial plateau cartilage (**Fig 4A**). Male mice exhibit worse articular damage and synovitis in this model, as we have thoroughly characterized^14^.

µCT imaging illustrates the formation of osteophytes and chondrophytes at the anterior meniscus, the anterior femur, and the posterior tibia (**Fig. 4B**), highly characteristic of this model. These can be contoured in 3D to derive quantitative osteophyte formation metrics, which we’ve shown to vary between male and female mice^14^. In addition to osteophyte formation, intra-synovial ossification may also be observed (**Fig. 4B**).

## Discussion

We present a comprehensive open-source protocol with associated quality control-enabling analysis tools and data to operationalize the murine ACLR model with a relatively low-cost, commercially available, low footprint, mobile mechanical testing system. Comparative mechanical data outputs demonstrate a high degree of parity with that of our conventional method using a high-cost ElectroForce instrument, and mechanical data in *n*=952 MoJO ACLRs demonstrates a very high degree of repeatability and success. Male and female mice exhibited the expected induction of knee hyperalgesia, synovial cell changes, osteophyte formation, and articular damage. Quantitative measures from these assessments were highly comparable to data collected with the conventional ElectroForce protocol, demonstrating a successful and accurate adaptation of our protocol from the ElectroForce system to the Univert S2 system.

### Advantages and Disadvantages of the ACLR Model

Benefits and drawbacks of ACLR as a preclinical model of PTOA are summarized in Table 1. As a noninvasive model with a clinically-relevant injury mechanism, ACLR is a uniquely well-suited model for studying the acute injury responses of joint tissues; in particular, the absence of a surgical incision to the synovium makes ACLR especially advantageous for study of synovial biology. The protocol is rapid, repeatable, and well-tolerated by mice, and produces rapid and progressive structural and pain-relevant changes. Conversely, the aggressive onset of articular destruction in this model, coupled with persistent joint instability, may make this a challenging model to implement for the assessment of long-term treatment strategies.

**Table 1.** Advantages and Disadvantages of ACLR compared to other preclinical models of PTOA.

| **Advantages** |  |
| --- | --- |
| Entirely noninvasive | - Not categorized as surgical procedure for IACUC<br>- Doesn't require sham procedures as control |
| Highly repeatable |  |
| Accurate induction of clinically-relevant joint injury | - Recapitulates motion profile of non-contact, sports-related ACL injury<sup>9,14</sup> |
| Rapid protocol | - ~5 mins per mouse |
| Good tolerability | - No weight loss<br>- No obvious signs of ill health or high pain |
| Straightforward, logical controls | - Contralateral and/or sham joints |
| Ability to assess early post-injury timepoints | - No confounding due to surgical approach |
| Useful for studying pain-correlated PTOA manifestations | - Synovitis, knee hyperalgesia, osteophyte formation |
| Useful for studying sex differences | - Observed sexual dimorphism in phenotypic presentation<sup>1</sup><br>- Female mice also exhibit PTOA development |
| **Disadvantages** |  |
| Rapid onset/progression of articular damage; high endpoint severity | - Scientists interested in studying milder articular cartilage and meniscus damage should consider the DMM or PMx models |
| Persistent joint instability | - May make implantation of chondral treatments challenging |
| Overlapping phases of post-injury inflammation and established PTOA | - May make treatment effects and timing difficult to determine or interpret |
| Concomitant meniscal damage/tears may be unwanted in specific studies | - Scientists interested in preserving the meniscus should consider the surgical ACL transection model |

### Controls: Contralateral limbs vs separate Sham mice

A major advantage of noninvasive PTOA models is the ability to use contralateral joints as internal controls. We recently demonstrated that in the ACLR model, contralateral joints do not exhibit any injury-induced joint damage, synovitis, osteophyte formation, or transcriptional changes in the synovium compared to separate uninjured Sham mice, within 28 days post-injury^14^. Flow cytometry data of the various synovial cell populations corroborates this further (data not shown, unpublished). The Sham procedure for the ACLR model involves a single anesthetic cycle and identical analgesia. As expected, Sham mice exhibit no nociceptive or tissue-level changes. Thus, these mice are effectively naïve, healthy controls, and both limbs may be utilized. While some groups may choose to perform Sham loading by administering only the preload and preconditioning segments of the protocol, these are not expected to induce any OA-relevant manifestations. Sham loading may nonetheless be important in mechanobiological studies in cartilage, meniscus, and bone, as we have previously measured very mild, acute increases in density of condylar bone in preload and preconditioning shams in an analogous rat model of ACLR^17^.

### Timepoints

Our prior published work in mice has shown early but detectable cartilage damage at 7d^8, 14^, concurrent with robust synovitis and the onset of synovial fibrosis. Other studies from our group and unpublished data have shown that monocyte/macrophage recruitment to the synovium is evident as early as 24-72 hrs post-injury^8^, with neutrophil influx happening as early as 8 hrs. Ectopic mineralization and osteophyte formation is evident at 10-14 days post-injury, but the earliest signs of chondrophyte formation are detectable as early as 7 days post-injury. Periarticular trabecular bone remodeling is highly dynamic and is evident by 7 days post-injury, but subchondral bone sclerosis is detectable after 14 days. By 28 days, we observe moderate PTOA severity in females and established/advanced severity in males^14^.

### Target Displacement Adjustment

Our mechanical data shows that in C57BL/6 mice aged 12-14 weeks, ACL rupture occurs at 0.9 – 1.3 mm in ∼90% of mice. We have chosen to target 1.5 mm to ensure that >99% of mice exhibit a complete ACL rupture, as our standard protocol does not allow re-loading of a failed rupture, and mice with failed ruptures are immediately euthanized. Thus, the additional 0.2 – 0.6 mm of post-rupture displacement is a safeguard against incomplete/failed rupture to minimize mouse utilization, consistent with the 3R principles. Some groups may wish to employ a lower target displacement of 1.25 or 1.3 mm, either when working with smaller mice or to avoid over-subluxation of the joint. Histological analysis demonstrates that in this model, anterior subluxation of the posterior meniscus can occur, which may drive greater articular damage.

It is likely possible to modulate the severity of this model, either by reducing post-rupture displacement below 1.5 mm in the setting of a complete rupture, or by decreasing displacement to induce subcritical injuries. By reducing loading, phenotypic manifestations such as articular damage and osteophytes may be able to be mitigated. While our group has not tested this in this specific model, other groups have employed subcritical cyclic or single-cycle loading to induce PTOA in mice^12, 13, 18, 19^. Further, low-level cyclic tibial compression was shown to exhibit therapeutic effects in DMM-induced OA^20^, while surgical restabilization of the murine joint following noninvasive ACLR was recently demonstrated, with promising evidence that restabilization can mitigate disease onset, particularly when coupled with joint unloading^21, 22^.

## Conclusion

Mouse models will continue to be an important tool for uncovering the causal disease mechanisms of OA and PTOA while also being a part of the development and translation of novel disease-modifying treatments. To facilitate the continued standardization of models with the goal of greater reproducibility across studies and sites, this study presented an open-source protocol and validation dataset of a mobile ACLR system.

## Acknowledgements

The authors thank Mr. Matt Brunsting, P.Eng., of CellScale for his extensive assistance and support in operationalizing our protocol on the Univert S2 system. We thank Dr. Aimee Colbath for kindly providing the ACLR mechanical data from her laboratory.

The RE-JOIN consortium consists of: Armen Akopian, Kyle Allen, Alejandro Almarza, Benjamin Arenkiel, Maryam Aslam, Basak Ayaz, Yangjin Bae, Bruna Balbino de Paula, Anita Bandrowski, Mario Danilo Boada, Jacqueline Boccanfuso, Jyl Boline, Dawen Cai, Dellina Lane Carpio, Robert Caudle, Racel Cela, Yong Chen, Rui Chen, Brian Constantinescu, Yenisel Cruz-Almeida, M. Franklin Dolwick, Chris Donnelly, Zelong Dou, Joshua Emrick, Malin Ernberg, Danielle Freburg-Hoffmeister, Jeremy Friedman, Spencer Fullam, Janak Gaire, Akash Gandhi, Terese Geraghty, Benjamin Goolsby, Stacey Greene, Nele Haelterman, Zhiguang Huo, Michael Iadarola, Shingo Ishihara, Sudhish Jayachandran, Zixue Jin, Alisa Johnson, Frank Ko, Zhao Lai, Brendan Lee, Yona Levites, Carolina Leynes, Jun Li, Martin Lotz, Lindsey Macpherson, Tristan Maerz, Camilla Majano, Anne-Marie Malfait, Maryann Martone, Simon Mears, Bella Mehta, Emilie Miley, Rachel Miller, Richard Miller, Michael Newton, Alia Obeidat, Soo Oh, Merissa Olmer, Dana Orange, Miguel Otero, Kevin Otto, Folly Patterson, Marlena Pela, Daniel Perez, Sienna Perry, Theodore Price, Hernan Prieto, Russell Ray, Dongjun Ren, Margarete Ribeiro Dasilva, Alexus Roberts, Elizabeth Ronan, Oscar Ruiz, Shad Smith, Mairobys Soccorro Gonzalez, Kaitlin Southern, Joshua Stover, Michael Strinden, Hannah Swahn, Evelyne Tantry, Sue Tappan, Cristal Villalba Silva, Airam Vivanco-Estella, Robin Vroman, Joost Wagenaar, Lai Wang, Kim Worley, Joshua Wythe, Jiansen Yan, and Julia Younis.

Research reported in this publication was supported by the National Institute of Arthritis and Musculoskeletal and Skin Diseases of the National Institutes of Health under Award Number P30 AR069620. The content is solely the responsibility of the authors and does not necessarily represent the official views of the National Institutes of Health.

## SUPPLEMENTAL INFORMATION

**Supplemental Figure 1.**
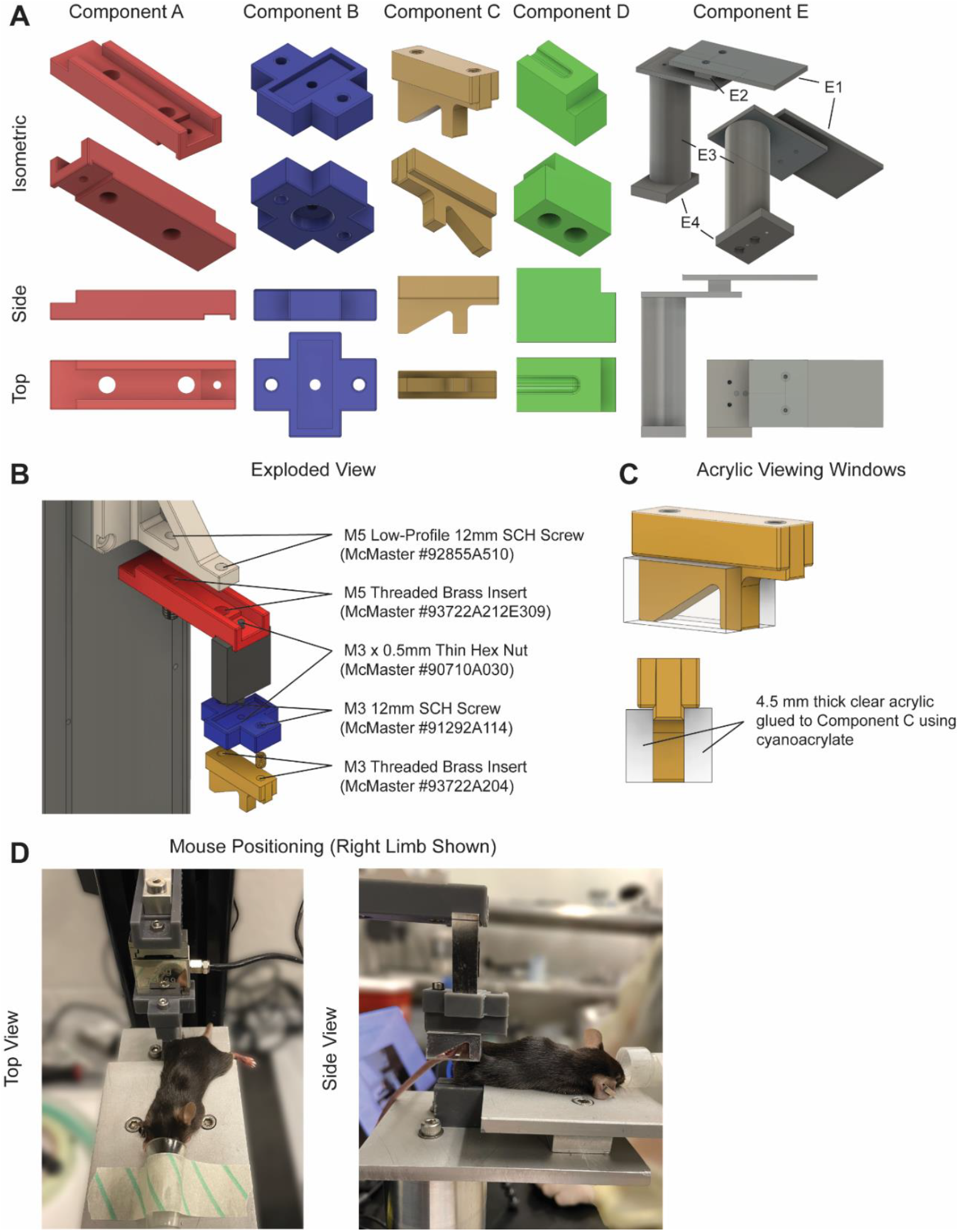
MoJO fixture components, assembly, and positioning. (A) Isometric, side, and top views of MoJO ACL rupture fixture components A-E. (B) Exploded view of upper fixture assembly. Fastener locations and part numbers are denoted. (C) Installation of acrylic viewing windows on component C (hindpaw fixture). (D) Top and side views of a mouse with right limb properly positioned within the ACL rupture fixture. Note that the head is in the anesthesia nosecone, the body is fully extended, the ankle and knee are fully seated at the front of their respective holders, and that the body is positioned such that the head, hip, knee, and ankle are approximately aligned. To achieve this positioning, the contralateral hip is externally-rotated to splay the contralateral limb on the animal bed, and the body and head are slightly rotated away from true prone positioning toward the loaded limb (in this case, to the right).

**Supplemental Figure 2.**
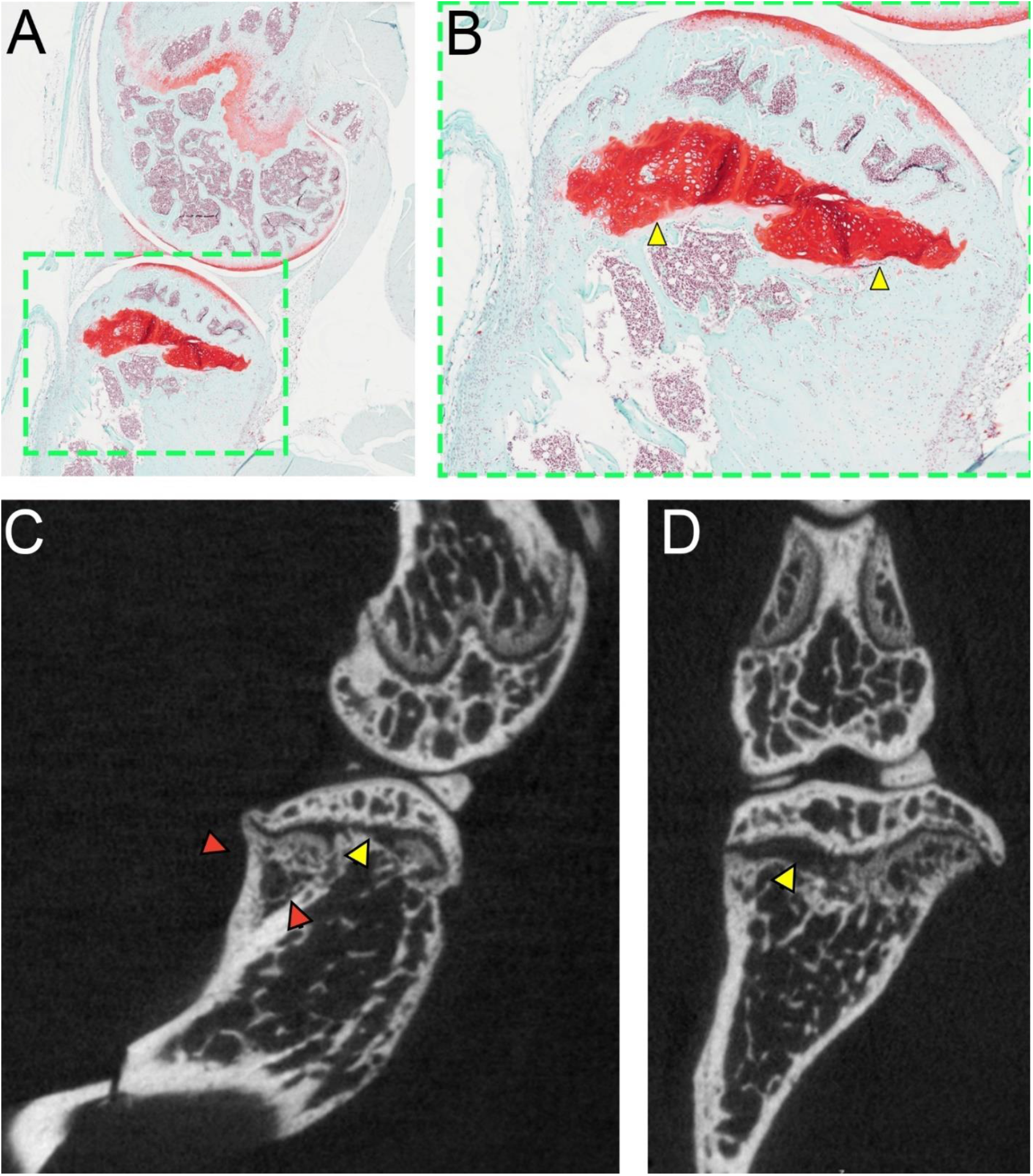
Example of a healed physeal rupture. (A) Sagittal Safranin-O-stained histological section of the medial joint compartment, including (B) a close-up image of the tibial plateau, demonstrating an abnormally large physis which has been filled in by growth plate cartilage (yellow arrows). (C) Sagittal and (D) coronal microCT sections demonstrate an enlarged tibial physis (yellow arrows), along with the presence of ectopic bone (red arrows) which appears to have grown in to support a posteriorly-shifted tibial epiphysis.

**Supplemental Figure 3.**
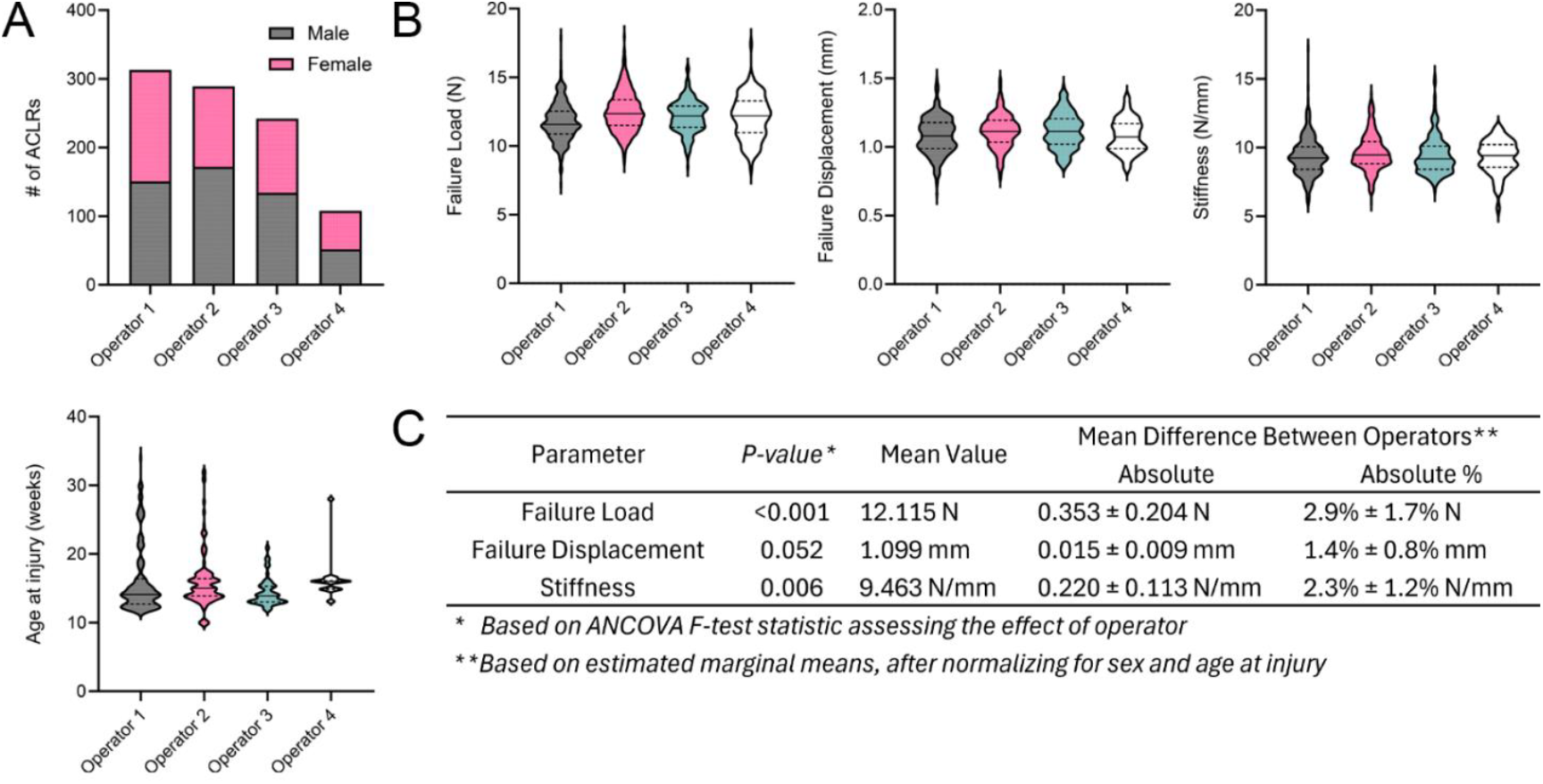
Reproducibility of MoJO ACL rupture outcomes between operators. (A) Distributions of sex and age at injury among 4 MoJO operators. (B) Mechanical parameters of ACL ruptures by operator, including failure load, failure displacement, and stiffness. (C) Statistical analysis of differences in mechanical outcomes between operators was assessed via ANCOVA, with operator as the between-subject factor and sex and age at injury as cofactors.

**Supplemental Figure 4.**
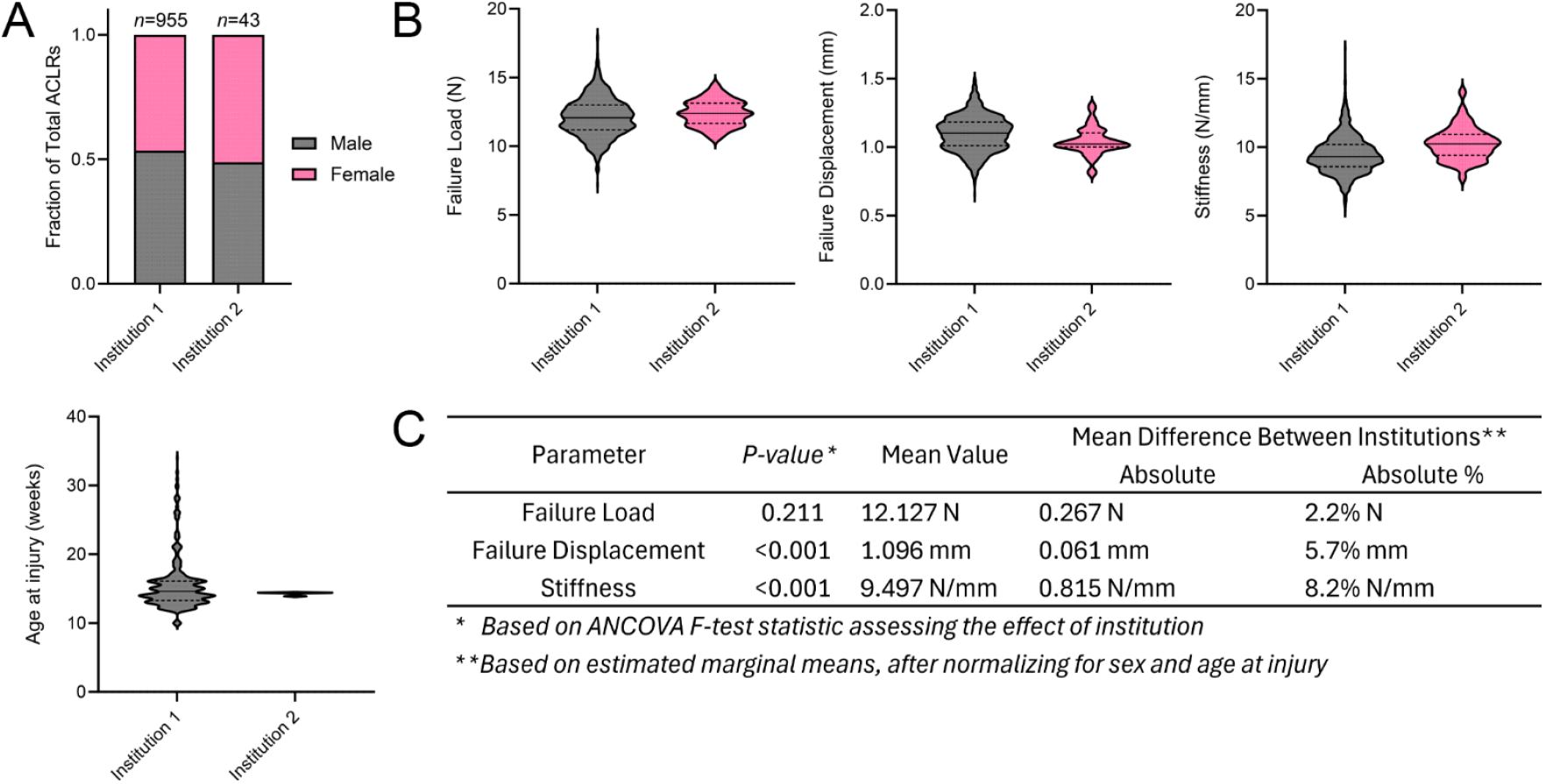
Reproducibility of MoJO ACL rupture outcomes between institutions. (A) Distributions of sex and age at injury among 2 institutions. (B) Mechanical parameters of ACL ruptures by institution, including failure load, failure displacement, and stiffness. (C) Statistical analysis of differences in mechanical outcomes between laboratories was assessed via ANCOVA, with institution as the between-subject factor and sex and age at injury as cofactors.

**Supplemental Methods 1** – *Additional Information on Fixture Design*

The most critical aspect of our fixture design is the alignment between the hindpaw fixture (C) and knee fixture (D), which controls knee flexion angle and hindlimb position to facilitate ACL rupture and minimize the risk of tibial fracture. The hindpaw fixture (C) is positioned slightly cranial to the knee fixture (D) to facilitate anterior femoral subluxation, which is required for loading of the ACL. The components are aligned such that the front of the knee sits 1.5 mm cranial to the front of the ankle. Importantly, to minimize mechanical moments that may induce fixture deflection and aberrant load cell readings, the approximate mechanical axis of the applied load traveling down the tibia is in-line with the center axis of the load cell. ***Critical: Any fixture modifications cannot alter the relative position between components C and D***.

If utilized with the CellScale Univert S2 and the associated ZMSA 2 kg load cell, all fixtures can be used without modification and as described in our design. If adapting to a different mechanical testing system, components **A** and **B** require machine-specific modifications. Alternatively, new adapters amenable for a given mechanical testing system may be designed to interface with components A and B. Similarly, if a different load cell besides the ZMSA 2 kg load cell (part of the CellScale Univert S2 system) is used, the load cell adapters (components A and B) will require modification, or new adapters that interface with components A and B can be made.

The cylinder (E4) is only needed to raise the animal bed into the stroke range of the Univert S2, and this component can be modified in length or eliminated entirely if utilizing a different mechanical testing system. It is highly preferable to raise the animal bed towards the crosshead using an extension such as the cylinder (E4) as opposed to placing an extension on the crosshead and component A, B, C assembly. The animal bed (E1) is large enough to accommodate mice of all sizes in addition to a standard anesthesia nose cone, which can be fastened to the bed using tape or a custom fixture. To enable visualization of hindpaw positioning within the hindpaw fixture (C), this part requires two transparent plastic platens to be glued on either side of component C. If machining is not available, these platens can be hand-cut using a sharp knife from 4.5 mm thick clear acrylic. The platens are then glued to component C using cyanoacrylate (**Suppl. Fig 1C**).

The knee fixture (D) contains a 3.5-mm wide by 1.5-mm deep trough that secures the knee in the medial-lateral and cranial directions. This geometry should fit mice of all sizes, but the trough may be widened for especially large mice, ensuring that the centerline of the trough does not move. Fillets are used to round all edges of this component to avoid skin injuries.

**Supplemental Methods 2** – *Diagnosis of Successful and Unsuccessful ACL Rupture Procedures*

During the first preload and during preconditioning cycles, the joint will exhibit viscoelastic creep. A gradual reduction in creep will be observed, with relatively stable load-displacement behavior achieved by the end of the second preload. This ensures the joint is fully seated and the ACL is fully engaged during the final compressive displacement, maximizing the rate of success and a full thickness rupture. During the final compressive 1.5-mm displacement, the load response of the joint prior to ACL rupture should be highly linear. If this is not the case, the viscous response of the ligament was not completely worked out during preloading. This can be occasionally observed in very large mice, or can reflect that the limb was not fully seated in the ankle and knee troughs at the start of the protocol, and this much of the “creep” observed simply represented the ankle and knee gradually slipping further into the fixture. Regardless, incomplete pre-injury engagement of the ACL does not necessarily guarantee an unsuccessful rupture – if the rupture displacement is characteristic of a successful ACL rupture, this still represents a successful procedure.

In our hands, the success rate of this procedure is incredibly high (99.0%), but unsuccessful procedures do occur. Representative load vs time plots of unsuccessful procedures are shown in **Fig. 1G**. Occasionally, the 1.5 mm displacement may prove insufficient to induce ACL rupture, with a higher risk in larger mice (>30g). In this case, no drop in load will be observed until the crosshead reverses direction, no audible “pop” will be heard, and no/minimal tibial subluxation will be observed. This is likely a consequence of improper seating of the hindlimb and high body fat.

Tibial fracture can also occur during this procedure and is a greater risk for smaller mice. Fractures typically occur in the midshaft, will produce a sharp audible sound of a different timbre than an ACL rupture, and will typically result in abrupt, *complete* unloading of the joint, without a substantial “catch” or re-loading. Suspected fractures can be assessed via manipulation of the tibia in traction, and via x-ray imaging. The mouse should be kept anesthetized throughout diagnosis, and euthanized *immediately* if a fracture has occurred. If a fracture is suspected but not confirmed, the mouse should be monitored during recovery and follow-up health checks for signs of abnormal gait and excessive pain, and follow-up x-rays can be taken as needed.

We have also observed instances of tensile rupture of the proximal tibial physis. This mode of failure is the most difficult to diagnose, but tends to produce a quieter, more gradually yielding sound. We have observed that the loading curve exhibits a more gradual, ductile yield and failure compared to the abrupt failure observed in an ACL rupture. A drop in load will occur, but rather than a sharp decrease in load followed by re-loading, a more gradual, “tumbling” unloading is observed as the physis yields and epiphysis displaces from the tibia. Suspected physeal dislocation can be assessed by ranging the knee joint, as displacement can result in either reduced range of motion or a distinct “crunchiness” arising from the epiphysis and metaphysis rubbing against one another as the physis shifts – range of motion can also be assessed during health checks the following day. X-rays can be acquired, but if the epiphysis settles back in its original position, this can be difficult to diagnose. Our group missed a tibial physis rupture and identified it during histological assessment, which shows cartilaginous healing tissue and a grossly widened physis (**Suppl. Fig 2A**). µCT images demonstrate substantial extracortical ectopic bone formation in the metaphysis underneath a shifted physis (**Suppl. Fig 2B**).

**Supplemental Methods 3** – *Calibration Spring Tuning*

The average stiffness of the mouse knee joint, i.e. ACL, in the linear region of Step 5 in this loading protocol is ∼10 N/mm. The preconditioning cycles are an important part of the protocol to ensure complete seating of the knee joint in the fixture while also preparing the joint for anterior subluxation. During the preconditioning cycles (Step 2), lower stiffnesses of ∼3-5 N/mm are expected. To assist in PID tuning and to perform protocol testing without mice, we recommend using steel compression springs with spring rates in this range. We have employed Product #9657K94 from McMaster-Carr (steel compression spring, 2.8” long, 0.625” outer diameter, 31 lbs/in spring rate, 54 lbs max load), and have found that tuning with this spring translates accurately to live mice. We recommend running the protocol on a test spring prior to each testing day, to confirm proper system response before loading live mice. We also collect longitudinal data on our spring tests to monitor system performance over time and ensure no drift occurs.

***Critical: Test each protocol or tuning parameter modification using steel springs, as recommended above, prior to using mice***.

**Supplemental Methods 4** – *Flow Cytometry of Synovial Cell Populations*

Synovia were digested using Type IV Collagenase, Liberase TM, and DNaseI for 30 mins, as previously described^15^. Cells were pre-blocked with Mouse TruStain FcX PLUS (Biolegend, clone S17011E) then stained with fluorescently conjugated antibodies: CD11b-BV510 (Biolegend, clone M1/70), CD146-BV605 (BD OptiBuild, clone ME-9F1), CD45-BV650 (Biolegend, clone 30-F11), CD31-PE/Cy7 (Biolegend, clone 390), F4/80-APC/R700 (BD Horizon, clone T45-2342), and CD3-APC/Fire750 (Biolegend, clone 17A2). Dead cells were excluded based on uptake of TO-PRO-3 iodide. Flow cytometry was performed on a BD Fortessa and data were analyzed using FlowJo software v10.

## References

1. R. Liukkonen, M. Vaajala, V.M. Mattila, and A. Reito, Prevalence of post-traumatic osteoarthritis after anterior cruciate ligament injury remains high despite advances in surgical techniques. Bone Joint J 105-B (11), 2023, 1140–1148. 10.1302/0301-620X.105B11.BJJ-2023-0058.R1.

2. N.A. Friel, and C.R. Chu, The role of ACL injury in the development of posttraumatic knee osteoarthritis. Clin Sports Med 32 (1), 2013, 1–12. 10.1016/j.csm.2012.08.017.

3. C. Blaker, E. Clarke, and C. Little, Pathology associations between different joint tissues are injury specific: a histopathological comparison of murine acl injury models. Osteoarthritis and Cartilage 28, 2020, S201-S202.

4. H. Urban, C. Blaker, C. Shu, E. Clarke, and C. Little, Synovial inflammation in anterior cruciate ligament injury knees in mice: surgical vs non-surgical models. Osteoarthritis and Cartilage 28, 2020, S213-S214.

5. B. Christiansen, M. Anderson, C. Lee, J. Williams, J. Yik, and D. Haudenschild, Musculoskeletal changes following non-invasive knee injury using a novel mouse model of post-traumatic osteoarthritis. Osteoarthritis and cartilage 20 (7), 2012, 773–782.

6. S.M. South, M.C. Marlin, P. Mehta-D’souza, T. Stephens, T. Conner, K.G. Burt, et al., Imaging mass cytometry reveals tissue-specific cellular immune phenotypes in the mouse knee following ACL injury. Osteoarthritis and Cartilage Open 5 (4), 2023, 100416.

7. C.L. Blaker, D.M. Ashton, N. Doran, C.B. Little, and E.C. Clarke, Sex-and injury-based differences in knee biomechanics in mouse models of post-traumatic osteoarthritis. Journal of Biomechanics 114, 2021, 110152.

8. P. Rzeczycki, C. Rasner, L. Lammlin, L. Junginger, S. Goldman, R. Bergman, et al., Cannabinoid Receptor Type 2 is Upregulated in Synovium following Joint Injury and Mediates Anti-Inflammatory Effects in Synovial Fibroblasts and Macrophages. Osteoarthritis and cartilage 29 (12), 2021, 1720.

9. T. Maerz, M.D. Kurdziel, A.A. Davidson, K.C. Baker, K. Anderson, and H.W. Matthew, Biomechanical characterization of a model of noninvasive, traumatic anterior cruciate ligament injury in the rat. Annals of biomedical engineering 43, 2015, 2467-2476.

10. T. Maerz, M. Newton, M. Kurdziel, P. Altman, K. Anderson, H. Matthew, et al., Articular cartilage degeneration following anterior cruciate ligament injury: a comparison of surgical transection and noninvasive rupture as preclinical models of post-traumatic osteoarthritis. Osteoarthritis and cartilage 24 (11), 2016, 1918–1927.

11. A. Ramme, M. Lendhey, J. Raya, T. Kirsch, and O. Kennedy, A novel rat model for subchondral microdamage in acute knee injury: a potential mechanism in post-traumatic osteoarthritis. Osteoarthritis and cartilage 24 (10), 2016, 1776–1785.

12. F.C. Ko, C. Dragomir, D.A. Plumb, S.R. Goldring, T.M. Wright, M.B. Goldring, et al., In vivo cyclic compression causes cartilage degeneration and subchondral bone changes in mouse tibiae. Arthritis & Rheumatism 65 (6), 2013, 1569–1578.

13. B. Poulet, R.W. Hamilton, S. Shefelbine, and A.A. Pitsillides, Characterizing a novel and adjustable noninvasive murine joint loading model. Arthritis & Rheumatism 63 (1), 2011, 137–147.

14. R.F. Bergman, L. Lammlin, L. Junginger, E. Farrell, S. Goldman, R. Darcy, et al., Sexual dimorphism of the synovial transcriptome underpins greater PTOA disease severity in male mice following joint injury. Osteoarthritis and Cartilage 32 (9), 2024, 1060–1073.

15. A.J. Knights, E.C. Farrell, O.M. Ellis, L. Lammlin, L.M. Junginger, P.M. Rzeczycki, et al., Synovial fibroblasts assume distinct functional identities and secrete R-spondin 2 in osteoarthritis. Annals of the rheumatic diseases 82 (2), 2023, 272–282.

16. A.J. Knights, E.C. Farrell, O.M. Ellis, M.J. Song, C.T. Appleton, and T. Maerz, Synovial macrophage diversity and activation of M-CSF signaling in post-traumatic osteoarthritis. eLife 12, 2025, RP93283. 10.7554/eLife.93283.

17. T. Maerz, M.D. Newton, M. Fleischer, S.E. Hartner, K. Gawronski, L. Junginger, et al., Traumatic joint injury induces acute catabolic bone turnover concurrent with articular cartilage damage in a rat model of posttraumatic osteoarthritis. Journal of Orthopaedic Research® 39 (9), 2021, 1965–1976.

18. C.L. Blaker, S. Zaki, C.B. Little, and E.C. Clarke, Long-term effect of a single subcritical knee injury: increasing the risk of anterior cruciate ligament rupture and osteoarthritis. The American Journal of Sports Medicine 49 (2), 2021, 391–403.

19. S. Ishihara, C. Blaker, R. Miller, E. Clarke, C. Little, and A.-M. Malfait, Critical and sub-critical load-induced knee injuries promote long term pain and impaired locomotion in mice. Osteoarthritis and Cartilage 26, 2018, S74.

20. D.T. Holyoak, C. Chlebek, M.J. Kim, T.M. Wright, M. Otero, and M.C. van der Meulen, Low-level cyclic tibial compression attenuates early osteoarthritis progression after joint injury in mice. Osteoarthritis and cartilage 27 (10), 2019, 1526–1536.

21. Y.-Y. Lin, E.H. Jbeily, P.M. Tjandra, M.C. Pride, M. Lopez-Torres, S.B. Elmankabadi, et al., Surgical restabilization reduces the progression of post-traumatic osteoarthritis initiated by ACL rupture in mice. Osteoarthritis and Cartilage 32 (8), 2024, 909–920.

22. Y.-Y. Lin, E. Jbeily, C. Lee, and B. Christiansen, Unloading Prior To Surgical Restabilization After Acl Rupture Slows Post-Traumatic Osteoarthritis Progression In Mice. Osteoarthritis and Cartilage 31, 2023, S205–S206.

